# Adversarial Attacks on Genotype Sequences

**DOI:** 10.1101/2022.11.07.515527

**Authors:** Daniel Mas Montserrat, Alexander G. Ioannidis

**Affiliations:** Stanford University

**Keywords:** Adversarial Attacks, Population Genetics, Genomics

## Abstract

Adversarial attacks can drastically change the output of a method by performing a small change on its input. While they can be a useful framework to analyze worst-case robustness, they can also be used by malicious agents to perform damage in machine learning-based applications. The proliferation of platforms that allow users to share their DNA sequences and phenotype information to enable association studies has led to an increase in large databases. Such open platforms are, however, vulnerable to malicious users uploading corrupted genetic sequence files that could damage downstream studies. Such studies commonly include steps involving the analysis of the genomic sequence’s structure using dimensionality reduction techniques and ancestry inference methods. In this paper we show how white-box gradient-based adversarial attacks can be used to corrupt the output of genomic analyses, and we explore different machine learning techniques to detect such manipulations.

## 1. INTRODUCTION

Open platforms such as [1, 2] allow users to upload their sequences and phenotype data (e.g. traits, medical records, ancestry information, etc). The available genomic sequences typically consist of single-nucleotide polymorphisms (SNPs), which are positions of the genome known to differ between individuals, and are often encoded as binary sequences with ‘0’ representing the common variant, and ‘1’ representing the alternative (minor) variant. These biobanks are beginning to be adopted in biomedical studies such as genome-wide association studies (GWAS) due to their free and easy access and their lack of privacy restrictions [3]. However, the free access to upload of sequences, make these platforms vulnerable to malicious users that can contribute corrupted genomes in order to infect downstream applications. For example, existing sequences could be modified or new sequences generated in order to disrupt the output and predictions of methods used in common bioinformatics pipelines, thus corrupting all downstream applications and analysis that rely on such predictive methods.

Bioinformatics pipelines used to perform association analyses (e.g. looking at correlations between genomic positions and phenotypes), phenotype prediction, or other biomedical applications, typically include dimensionality reduction techniques and clustering methods to both provide visualization tools and to extract lower dimensional representations of genomes which are then used as covariates in downstream analyses. Some common techniques include Principal Component Analysis (PCA), soft clustering such as Neural ADMIXTURE (N. ADM) [4], and ancestry deconvolution (also refered to as local ancestry inference) such as LAI-Net (LN) [5] and SALAI-Net [6], which assign a population label to each region of the chromosome sequence. As many of these techniques provide available code and pretrained models, they are vulnerable to white-box adversarial attacks. In this paper we introduce a technique to perform an adversarial attack that, by changing a small percentage of the input sequence, severely damages the output of these above methods.

Neural network-based techniques, which can be highly vulnerable to adversarial attacks, are beginning to be adopted within genomic pipelines for association analysis, including networks for variant calling [7], population structure inference [4, 5, 6], phenotype prediction [8], or neural networkbased GWAS [9]. As neural networks become more common in such pipelines, their robustness becomes increasingly important, and adversarial attacks provide a framework to evaluate their stability under perturbations of the input sequence. In fact, even if no malicious agents generating attacks are present, the corruption of segments of genomic sequences can happen during the sequencing process or during the preprocessing of the data. In such scenario, adversarial attack techniques provide a practical mechanism to perform worst-case robustness analysis.

Many adversarial attacks have been recently proposed [10], including white-box [11, 12] and black-box [13] attacks to perturb a method’s prediction during test time and poisoning attacks [14] to corrupt the training data. Likewise, many defense mechanisms have been proposed, such as adversarial training [15], data pre-processing [16], or manipulation detection [17]. While most of this previous work has been applied to images, video, text, or audio, genomic sequences still remain largely unexplored. To our knowledge this is the first work to apply adversarial perturbations within genotype sequences (SNPs).

## 2. ADVERSARIAL MANIPULATION

### 2.1. Adversarial Mutations

We introduce an adversarial attack to change the prediction of commonly used genotype classification and segmentation methods (i.e. global and local ancestry inference), while minimally modifying the input sequences. Note that common adversarial attacks add a small adversarial perturbation ε to the input *x*, obtaining an adversarial input *x*^*^ = *x* + *ϵ*, with an ε small or below a threshold under some norm. In our scenario, the input *x* ∈ {0, 1}^*d*^ is a *d*-dimensional binary sequence of SNPs, and we want the generated adversarial sequence to still be a *d*-dimensional binary sequence. In order to perform the adversarial attack, we use a *d*-dimensional binary ‘mutation mask’ *δ* ∈ {0, 1}^*d*^ that indicates which positions of the DNA sequence need to be changed (from a 0 to a 1 or vice versa). Hence, the adversarial (mutated) sequence, *x*\* can be represented as:

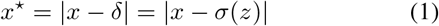

where *x*^*^, *x*, *δ* ∈ {0, 1}^*d*^ are the adversarial, input, and mutation sequence respectively. |·| is the element-wise absolute value function, *σ*(·) is a differentiable binarization layer, and 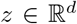 is a latent vector that parametrizes the mutation mask *δ* = *σ*(*z*). By parematrizing the mutation mask as a latent sequence through a differentiable binarization layer, we ensure that the mutation mask is always binary, while being able to backpropagate the gradients to update the latent sequence (and in turn the mutation mask). In this work we make use of the binarization layer proposed in [18], which proved successful on generating SNP sequences in [19].

The adversarial mutation sequence is obtained by solving the following optimization problem, based on the C&W [12] adversarial attack:

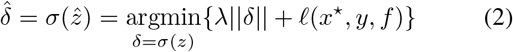

where ║*δ*║ is the L1 norm of the mutation mask, which is equivalent to the number of DNA positions that are changed, *ℓ*(·) is a loss function that evaluates how good the prediction of a given classifier or segmentation method *f*(·) is, and *λ* is a scaling hyperparameter to change the balance of the effect of the L1 norm. We use the same loss function as in the C&W attack:

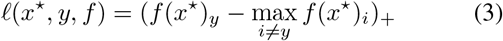

where (·)_+_ = max(·, 0), *y* is the true label of the input sample *x, f*(*x*^*^)_*y*_ is the value of the logit (or probability) for the true category y when using *x*^*^ as input, and max_*i*≠*y*_ *f*(*x*^*^)_*i*_ is the largest logit (or probability) different than the one assigned to the true category.

In practice, we obtain 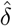 by iteratively updating *z* with an optimization algorithm that minimizes *L*(*x, z, f, y*) = *λ*║*σ*(*z*)║ + *ℓ*(|*x* – *σ*(*z*)|, *y, f*). In our experiments we make use of the Adam optimizer for 20 iterations with a learning rate of 0.05 and a lambda of *λ* = 1.0 for LAI-Net and *λ* = 3.0 for N. ADM.

### 2.2. Adversarial Robustness Analysis

After the mutation mask 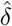 has been predicted for a given input sequence x, multiple additional mutation masks are generated by mixing 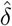 with a random (non-adversarial) binary sequence *δ_r_*. In practice, *δ_r_* is constructed such that 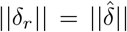 by performing a random permutation of 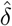. New mutation masks *δ_α,β_* are generated by randomly selecting elements from *δ_r_* and 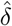:

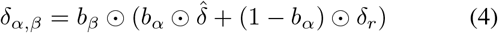

with *δ_r_, δ_α_, β, b_α_, b_β_* ∈ {0, 1}^*d*^ and ⊙ as the element-wise multiplication. *b_α_* ~ Bernoulli(**1***α*) and *b_β_* ~ Bernoulli(**1***β*) are d-dimensional binary masks where each dimension follows an independent Bernoulli distribution with parameter *α* and *β* respectively, and **1** is a d-dimensional vector of ones. *b_α_* is a binary mask that selects if *δ_α,β_* contains a mutation from the adversarial mutation sequence 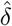 or the non-adversarial mutation sequence *δ_r_*. *b_β_* is a binary mask that selects how many elements from the mutation sequence are active (i.e. non-zero). With *α* = 1 and *β* = 1, we recover the adversarial mutation sequence 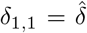, with *α* = 0 and *β* = 1, the random sequence *δ*_0,1_ = *δ_r_*, and with *β* = 0, *δ*_*α*,1_ = 0, and therefore *x*^*^ = *x*. Note that we can’t do simple linear interpolation as the resulting sequence could contain non-binary values. We use *δ_α,β_* to evaluate the robustness of the classification and segmentation methods under perturbations at the inputs sequences. In order to do so, we compute the classification (segmentation) accuracy under different values of *α* and *β*:

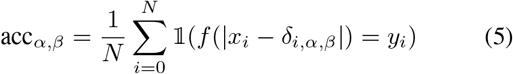

where 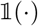 is the indicator function and *δ_i,α,β_* is the mutation mask *δ_α,β_* for the *i*th sample *x_i_* with label *y_i_*. acc_*α*, 0_ is the accuracy without any mutation, acc_1,1_ is the accuracy under adversarial mutations, and acc_0,1_ is the accuracy under random mutations.

### 2.3. Experimental Results

We use human whole genome sequences of chromosome 20 from three publicly available datasets including the 1000 Genomes Project (1kg) [20], the Simons Genome Diversity Project (SGDP) [21] and the Human Genome Diversity Project (HGDP) [22]. We have a total of 720 sequences with 516,000 SNPs for each population group. We obtain a balanced dataset by up-sampling the underrepresented populations using the simulation procedure adopted in [5]. The dataset is grouped into 8 population clusters: African (AFR), African Hunther-Gatherers (AHG), European (EUR), East Asian (EAS), South Asian (SAS), West Asian (WAS), Oceanian (OCE), and Native American (AMR).

We train LAI-Net [5] for sequence segmentation (local ancestry inference) and Neural ADMIXTURE [4] for sequence classification (global ancestry inference) with the previous dataset and evaluate their robustness under adversarial perturbations by computing *acc_α,β_* for multiple values of *α* and *β*. Figure 1 shows the accuracy as a function of percentage of manipulated SNPs and the amount of adversarial perturbation *α*. We can see that by changing a very small percentage of genomic positions, the accuracy drops from almost 100% to below 10%. Furthermore, we can see that if the perturbation is random and not adversarial (light colors), the accuracy is untouched. These results show that the adversarial attacks are successful, and while the methods are robust to small random perturbations, completely fail in the presence of adversarial attacks.

**Fig. 1.**
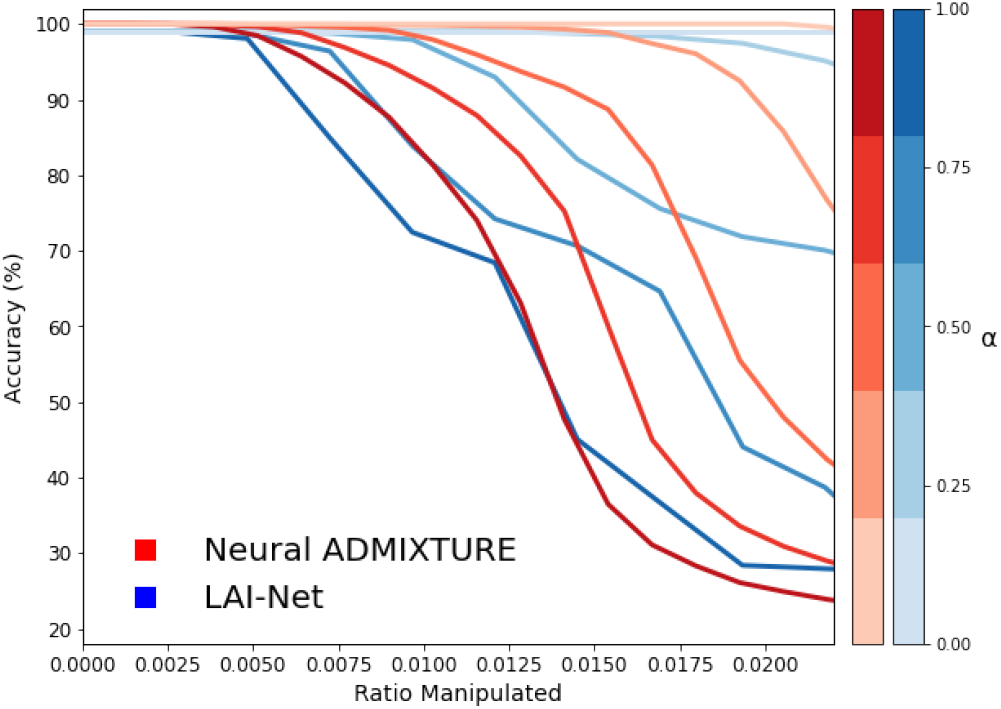
LAI-Net (Blue) and N. ADM (Red) accuracy under adversarial perturbations. Dark to light color indicates high to low *α*. Percentage manipulated is changed by varying *β*.

## 3. ADVERSARIAL SEQUENCE GENERATION

### 3.1. Targeted Population Generation

Synthetic adversarial sequences that produce a desired prespecified output from genetic processing methods are generated through backpropagation with a targeted loss function. Specifically, we show that a set of SNP sequences can be generated to provide the desired output of different techniques including PCA, Fast k-NN (k-NN applied to random projections of the data), LAI-Net, and N. ADM. As an example, we generate of 50 sequences with 516k SNPs with PCA projections that track European individuals, but that have semisupervised clustering outputs from Neural ADMIXTURE indicating they are instead Oceanian samples, are meanwhile classified by fast k-NN as African Hunter-Gatherers, and have LAI-Net classifying them as Native American. In order to do so, we generate a dataset 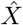 of SNP sequences by using a set of targeted loss functions:

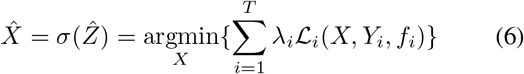

where 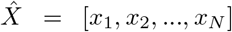 is a dataset with *N d*-dimensional binary sequences *x_i_* ∈ {0, 1}^*d*^, parematrized by 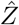. 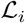 is a loss function applied for the task *i* out of *T* tasks. Each task makes use of a model *f_i_* and has a target label *Y_i_*. The loss for each task has a weight *λ_i_* assigned. In this paper we make use 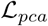 for dimensionality reduction, 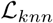 for k-NN classification, and 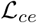 for LAI-Net and N. ADM classification. 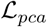 is as follows:

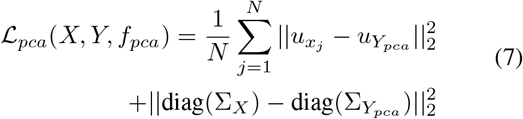

where *u_x_j__, u_pca_*, diag(Σ_*X*_), 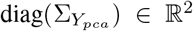. *u_x_j__* = *W_pca_*(*x_j_* – *b_pca_*) are the two first principal component coordinates of sample *x_j_*, and *W_pca_* and *b_pca_* the PCA projection matrix and the centering vector respectively, computed on the real sequences. 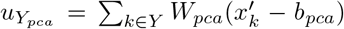 is the average of the two first principal component coordinates of the training real samples 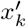 of the *Y_pca_* target category (in our example Europeans). diag(Σ_*X*_) and diag(Σ*_Y_pca__*) are the variance of the simulated and real data after projection to the PCA. This loss enforces that the projected PCA of the fake samples is close to the center of the specified cluster (e.g. Europeans), and that the variance of the projected points from fake and real sequences are similar. 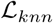 is as follows:

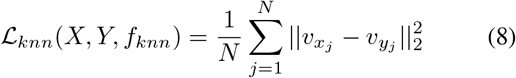

where *v_x_j__* = *W_knn_x_j_* is a random projection of the *j*th generated sequence *x_j_*, with *W_knn_* sampled from a Gaussian distribution. 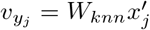 is a random projection of the *j*th real sequence 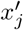 from the population group with label *y_j_* = *Y_knn_*. This loss enforces that the random projection of each simulated sequence is similar to the random projection of a real sequence from the target population *Y_knn_*. The loss used for LAI-Net and Neural ADMIXTURE, 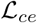, is the regular cross entropy (CE) loss.

### 3.2. Experimental Results

Figure 2 shows the result of multiple methods on real, random, and adversarial sequences. The random sequences are obtained by sampling from a Bernoulli distribution where each dimension has a probability equivalent to the empirical frequency of each respective SNP position. Adversarial sequences (red dots) are projected on top of European samples (green dots), making them visually indistinguishable from real sequences in a PCA plot. The top row shows the percentage of predictions for each cluster when testing European sequences are used as input, showing that the 3 methods correctly classify them. If random sequences are used as input (middle row), Fast k-NN and N. ADM provide random outputs, while LAI-Net classify them as OCE, which appears as a distinct cluster in PCA space. When the adversarial sequences are used as input (bottom row), each method outputs the category specified as target label (EUR for PCA, AHG for k-NN, AMR for LAI-Net, and OCE for N. ADM), showing that the target generation of sequences has been successful. These results provide the following insights: (1) synthetic fake sequences can lead to realistic-looking outputs from bioinformatics methods, (2) the sequences can be generated in such a way that any desired output can be obtained, leading to highly inconsistent (or consistent) outputs between different techniques, (3) commonly used dimensionality reduction and classification techniques can be easily fooled, showing that structure and scientific insight found using such techniques in genetic analyses should be handled with caution.

**Fig. 2.**
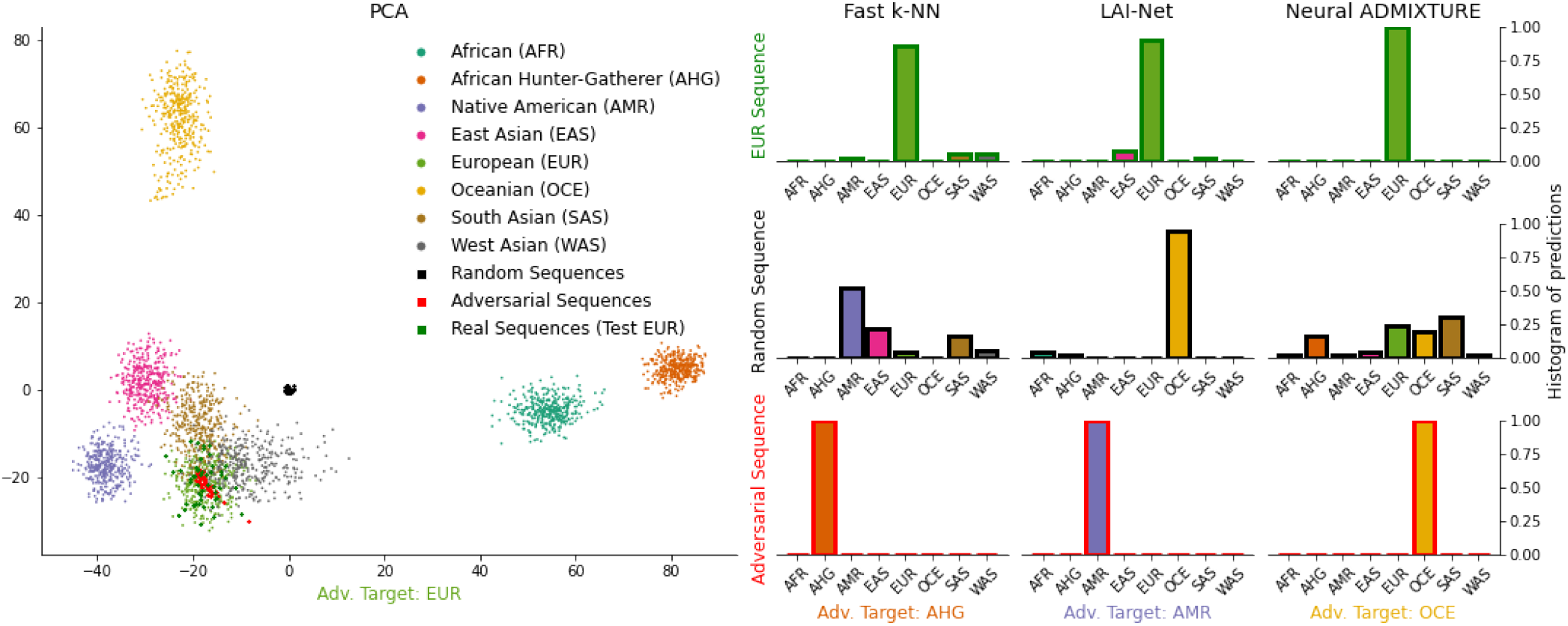
PCA, k-NN, LAI-Net, and Neural ADMIXTURE results from real (train and test), random, and adversarial sequences.

## 4. ADVERSARIAL ATTACK DETECTION

We explore the use of supervised learning classifiers to discriminate between real and adversarial fake sequences. We explore common techniques applied on genetic data: logistic regression, k-Nearest Neighbor, XGBoost, and a Multi-Layer Perceptron (MLP). A set of classifiers are trained to detect the adversarial manipulation applied to LAI-Net (LN) and N. ADM introduced in section 2, and another is trained to detect the adversarial synthetic (A.S.) generated sequences introduced in section 3. A total of 1440 real samples, 1440 manipulated and 50 generated generated sequences, are used to train and evaluate the methods with a 50-50 train-test split. Table 1 shows that most of the classifiers are able to accurately detect the manipulations and distinguish between real and fake sequences, indicating that such adversarial attacks leave distinctive patterns.

**Table 1.**
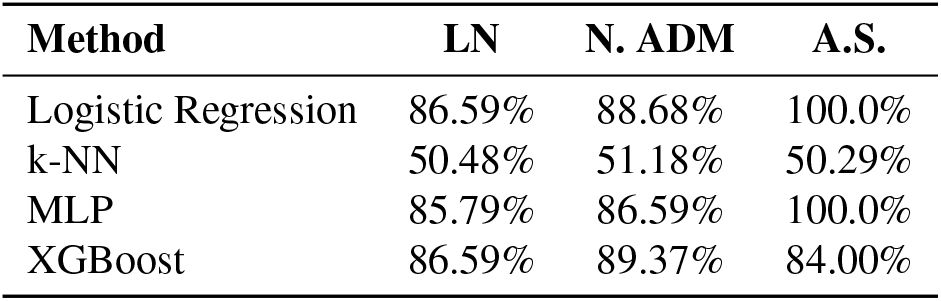
Manipulation detection balanced accuracy.

## 5. CONCLUSIONS

Adversarial attacks applied to genotypes have remained unexplored. Here we introduce, to the best of our knowledge, the first use of adversarial attacks on genomic sequences and showcase two different attacks, including sequence perturbation and sequence synthesis, applied to dimensionality reduction, classification, and segmentation techniques. While these attacks are effective, we show that they are easily detected by manipulation detection techniques.

